# Placozoa and Cnidaria are sister taxa

**DOI:** 10.1101/200972

**Authors:** Christopher E. Laumer, Harald Gruber-Vodicka, Michael G. Hadfield, Vicki B. Pearse, Ana Riesgo, John C. Marioni, Gonzalo Giribet

## Abstract

The phylogenetic placement of the morphologically simple placozoans is crucial to understanding the evolution of complex animal traits. Here, we examine the influence of adding new genomes from placozoans to a large dataset designed to study the deepest splits in the animal phylogeny. Using site-heterogeneous substitution models, we show that it is possible to obtain strong support, in both amino acid and reduced-alphabet matrices, for either a sister-group relationship between Cnidaria and Placozoa, or for Cnidaria and Bilateria (=Planulozoa), also seen in most published work to date, depending on the orthologues selected to construct the matrix. We demonstrate that a majority of genes show evidence of compositional heterogeneity, and that the support for Planulozoa can be assigned to this source of systematic error. In interpreting this placozoan-cnidarian clade, we caution against a peremptory reading of placozoans as secondarily reduced forms of little relevance to broader discussions of early animal evolution.

## Introduction

The discovery^1^ and mid-20^th^ century rediscovery^2^ of the enigmatic, amoeba-like placozoan *Trichoplax adhaerens* did much to ignite the imagination of zoologists interested in early animal evolution^3^. As microscopic animals adapted to extracellular grazing on the biofilms over which they creep^4^, placozoans have a simple anatomy suited to exploit passive diffusion for many physiological needs, with only six morphological cell types discernible even to intensive scrutiny^5, 6^, and have no conventional muscular, digestive, or nervous systems, yet show tightly-coordinated behavior^7, 8^. They proliferate through fission and somatic growth. Evidence for sexual reproduction remains elusive, despite genetic evidence of recombination^9^ and descriptions of early abortive embryogenesis^10, 11^, with the possibility that sexual phases of the life cycle may occur only under poorly understood field conditions^12, 13^

Given their simple, puzzling morphology and dearth of embryological clues, molecular data are crucial in placing placozoans phylogenetically. The position of Placozoa in the animal tree proved recalcitrant to early standard-marker analyses^14–16^, although this paradigm did reveal a large degree of molecular diversity in placozoan isolates from around the globe, clearly indicating the existence of many cryptic species^12, 17, 18^ with up to 27% genetic distance in *16S rRNA* alignments^19^. An apparent answer to the question of placozoan affinities was provided by analysis of a nuclear genome assembly^9^, which strongly supported a position as the sister group of a clade of Cnidaria+Bilateria (sometimes called Planulozoa). However, this effort also revealed a surprisingly bilaterian-like^20^ developmental gene toolkit in placozoans, a paradox for such a simple animal.

As metazoan phylogenetics has pressed onward into the genomic era, perhaps the largest controversy has been the debate over the identity of the sister group to the remaining metazoans, traditionally thought to be Porifera, but considered to be Ctenophora by Dunn et al.^21^ and subsequently by additional studies^22–26^. Others have suggested that this result arises from inadequate taxon sampling, flawed matrix husbandry, and/or use of poorly fitting substitution models^27–31^. A third view has emphasized that using different sets of genes can lead to different conclusions, with only a small number sometimes sufficient to drive one result or another^32, 33^. This controversy, regardless of its eventual resolution, has spurred serious contemplation of possibly independent origins of several hallmark traits such as striated muscles, digestive systems, and nervous systems^23, 34–39^.

Driven by this controversy, new genomic and transcriptomic data from sponges, ctenophores, and metazoan outgroups have accrued, while new sequences and analyses focusing on the position of Placozoa have been slow to emerge. Here, we provide a novel test of the phylogenetic position of placozoans, adding draft genomes from three putative species that span the root of this clade’s known diversity^17^, and critically assessing the role of systematic error in placing of these enigmatic organisms.

## Results and Discussion

Orthology assignment on sets of predicted proteomes derived from 59 genome and transcriptome assemblies yielded 4,294 gene trees with at least 20 sequences each, sampling all 5 major metazoan clades and outgroups, from which we obtained 1,388 well-aligned orthologues. Within this set, individual maximum-likelihood (ML) gene trees were constructed, and a set of 430 most-informative orthologues were selected on the basis of tree-likeness scores^40^. This yielded an amino-acid matrix of 73,547 residues with 37.55% gaps or missing data, with an average of 371.92 and 332.75 orthologues represented for Cnidaria and Placozoa, respectively (with a maximum of 383 orthologues present for the newly sequenced placozoan H4 clade representative; Figure 1).

**Figure 1.**
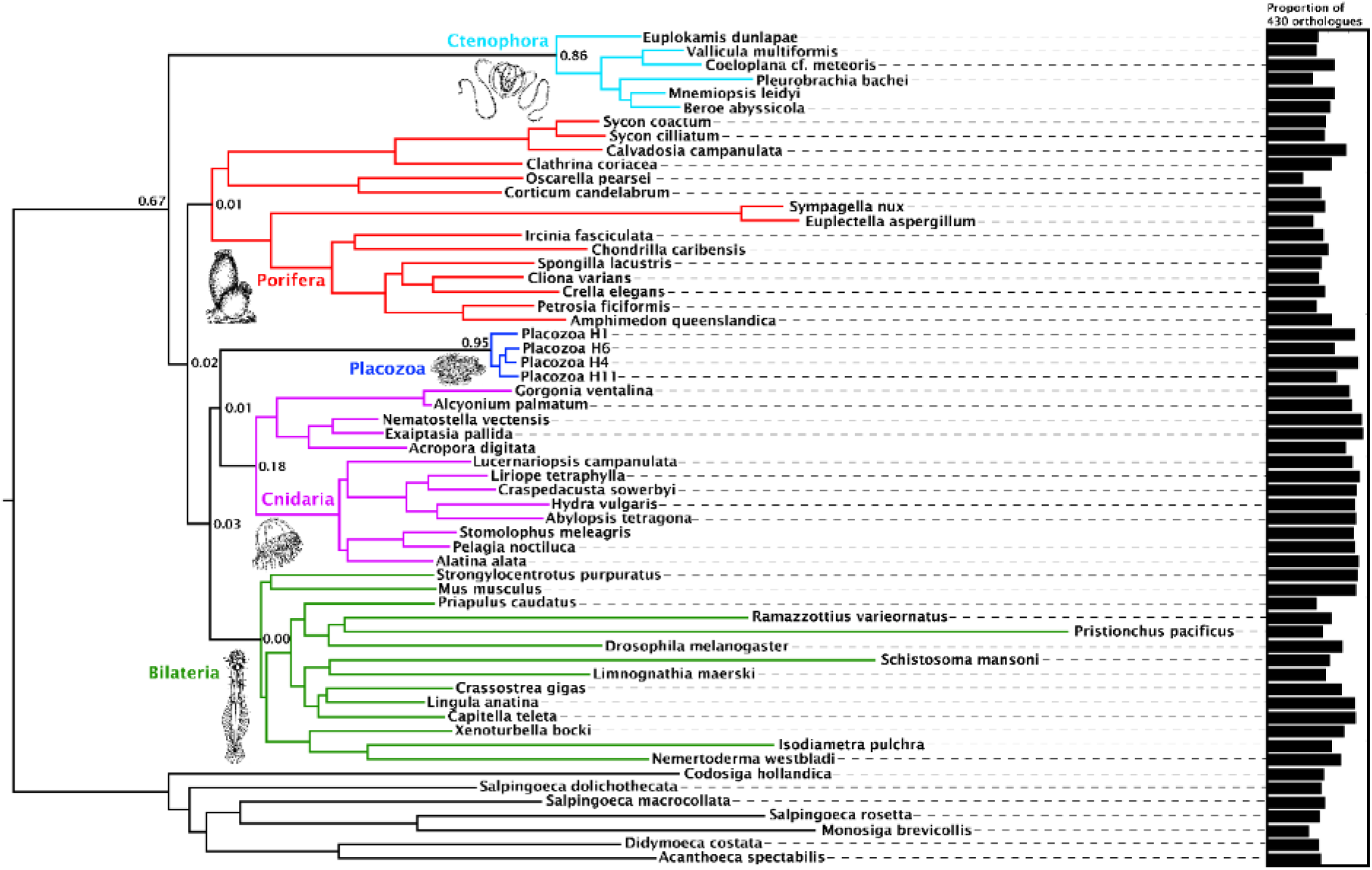
– Consensus phylogram showing deep metazoan interrelationships under Bayesian phylogenetic inference of the 430-orthologue amino acid matrix, using the CAT+GTR+Г4 mixture model. All nodes received full posterior probability. Numerical annotations of given nodes represent Extended Quadripartition Internode Certainty (EQP-IC) scores, describing among-gene-tree agreement for both the monophyly of the 5 major metazoan clades and the given relationships between them in this reference tree. A bar chart on the right depicts the proportion of the total orthologue set each terminal taxon is represented by in the concatenated matrix. ‘Placozoa H1’ in this and all other figures refers to the GRELL isolate sequenced in Srivastava et al 2008, which has there and elsewhere been referred to as *Trichoplax adhaerens*, despite the absence of type material linking this name to any modern isolate. Line drawings of clade representatives are taken from the BIODIDAC database (http://biodidac.bio.uottawa.ca/).

Our Bayesian analyses of this matrix place Cnidaria and Placozoa as sister groups with full posterior probability under the general site-heterogeneous CAT+GTR+Г4 model (Figure 1). Under ML inference with a profile mixture model^41^ (Figure 1 – Figure Supplement 1), we again recover Cnidaria+Placozoa, albeit with more marginal resampling support. Both Bayesian and ML analyses show little internal branch diversity within Placozoa. Accordingly, deleting all newly-added placozoan genomes from our analysis has no effect on topology and only a marginal effect on support in ML analysis (Figure 1 – Supplemental Figure 2). Quartet-based concordance analyses^42^ show no evidence of strong phylogenetic conflicts among ML gene trees in this 430-gene set (Figure 1), although internode certainty metrics are close to 0 for many key clades including Cnidaria+Placozoa, indicating that support for some ancient relationships may be masked by gene-tree estimation errors, emerging only in combined analysis^43^.

Compositional heterogeneity of amino-acid frequencies along the tree is a source of phylogenetic error not modelled by even complex site-heterogeneous substitution models such as CAT+GTR^44–47^. Furthermore, previous analyses^32^ have shown that placozoans and choanoflagellates in particular, both of which taxa our matrix samples intensively, deviate strongly from the mean amino-acid composition of Metazoa, perhaps as a result of genomic GC content discrepancies. As a measure to at least partially ameliorate such nonstationary substitution, we recoded the amino-acid matrix into the 6 “Dayhoff” categories, a common strategy previously shown to reduce the effect of compositional variation among taxa, albeit the Dayhoff-6 groups represent only one of many plausible recoding strategies, all of which sacrifice information^31, 48–50^. Analysis of this recoded matrix under the CAT+GTR model again recovered full support (pp=1) for Cnidaria+Placozoa (Figure 2). Indeed, under Dayhoff-6 recoding, the only major change is in the relative positions of Ctenophora and Porifera, with the latter here constituting the sister group to all other animals with full support. Similar recoding-driven effects on relative positions of Porifera and Ctenophora have also been seen in other recent work^31^.

**Figure 2.**
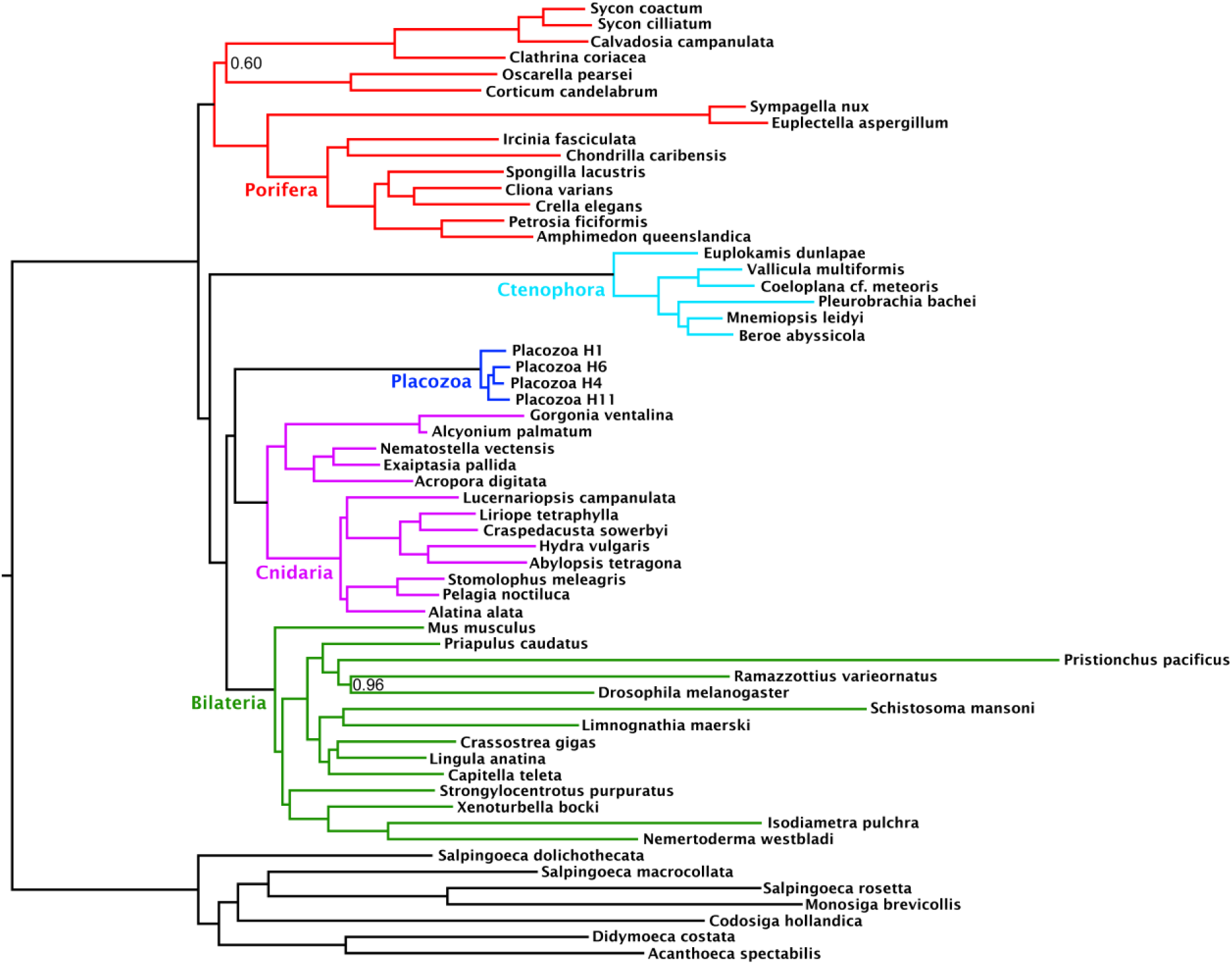
– Consensus phylogram under Bayesian phylogenetic inference under the CAT+GTR+Г4 mixture model, on the 430-orthologue concatenated amino acid matrix, recoded into 6 Dayhoff groups. Nodes annotated with posterior probability; unannotated nodes received full support.

Many research groups, using good taxon sampling and genome-scale datasets, and even recently including data from a new divergent placozoan species^26, 31, 51^, have consistently reported strong support for Planulozoa under the CAT+GTR model. Indeed, when we construct a supermatrix from our predicted peptide catalogues using a different strategy, relying on complete sequences of 303 pan-eukaryote “Benchmarking Universal Single-Copy Orthologs” (BUSCOs) ^52^, we also see full support in a CAT+GTR+Г analysis for Planulozoa, in both amino-acid (Figure 3a) and Dayhoff-6 recoded alphabets (Figure 3b). Which phylogeny is correct, and what process drives support for the incorrect topology? Posterior predictive tests, which compare the observed among-taxon usage of amino-acid frequencies to expected distributions simulated using the sampled posterior distribution and a single composition vector, may provide insight^46^. Both the initial 430-gene matrix and the 303-gene BUSCO matrix fail these tests, but the BUSCO matrix fails it more profoundly, with z-scores (measuring mean-squared across-taxon heterogeneity) scoring in the range of 330-340, in contrast to the range of 176-187 seen in the 430-gene matrix. Furthermore, inspecting z-scores for individual taxa in representative chains from both matrices shows that a large amount of this global difference in z-scores can be attributed to placozoans, with additional contributions from choanoflagellates and select isolated representatives of other clades (Figure 3C).

**Figure 3.**
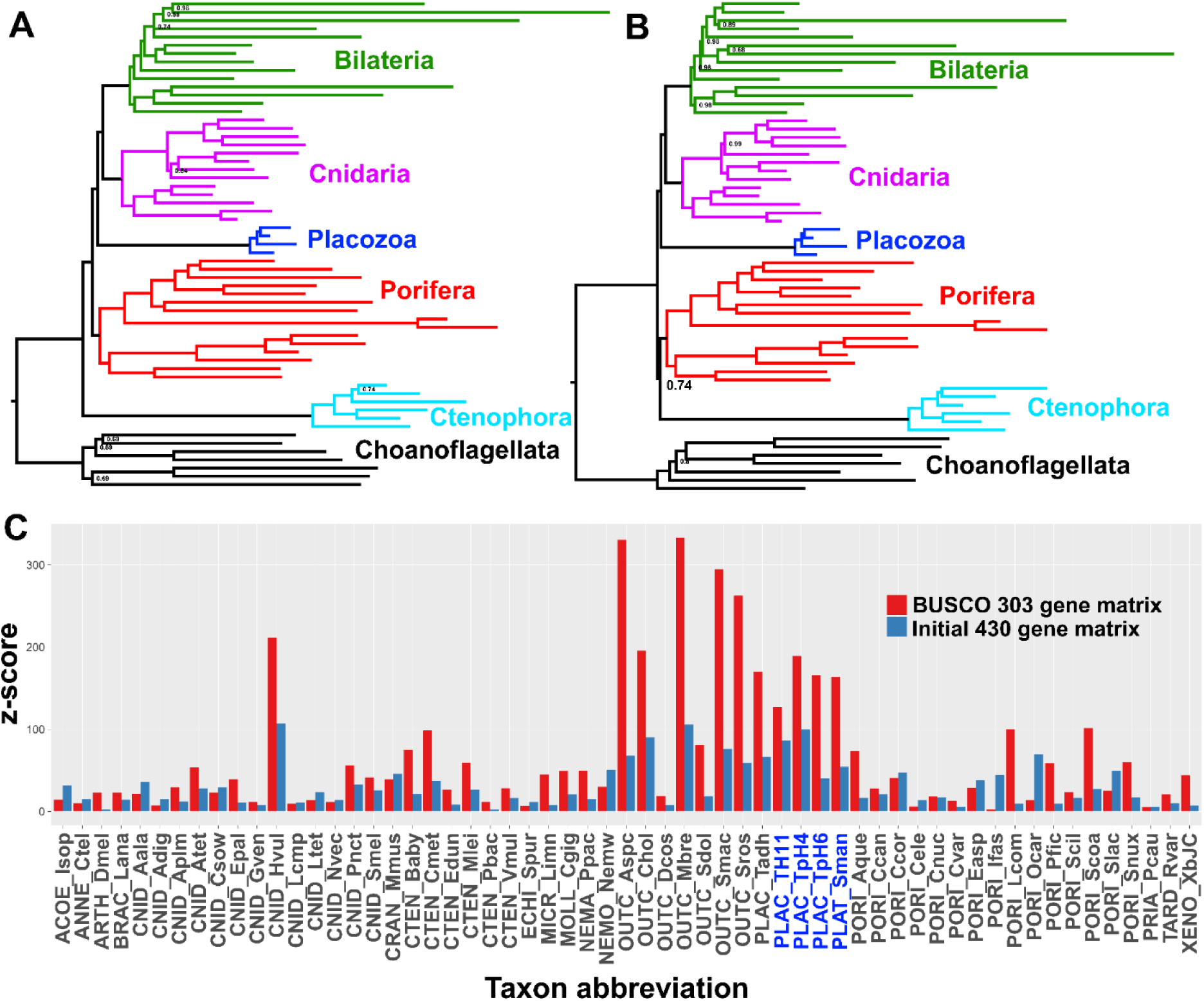
– Posterior consensus trees from CAT+GTR+Г4 mixture model analysis of a 94,444 amino acid supermatrix derived from the 303 single-copy conserved eukaryotic BUSCO orthologs, analysed in A.) amino acid space or B.) the Dayhoff-6 reduced alphabet space. Nodal support values comprise posterior probabilities; nodes with full support not annotated. Taxon colourings as in previous Figures. C.) Plot of z-scores (summed absolute distance between taxon-specific and global empirical frequencies) from representative posterior predictive tests of amino acid compositional bias, from both the BUSCO 303-orthologue matrix (red) and the initial 430-orthologue matrix (blue). Placozoan taxon abbreviations are shown in blue font.

As a final measure to describe the influence of compositional heterogeneity in this dataset, we applied a null-simulation test for compositional bias to each alignment in our set of 1,388 orthologues. This test, which compares the real data to a null distribution of amino-acid frequencies simulated along assumed gene trees with a substitution model using a single composition vector, is less prone to Type II errors than the more conventional Χ^2^ test^45^. Remarkably, at a conservative significance threshold of α=0.10, the majority (764 genes or ∼55%) of this gene set is identified as compositionally biased by this test, highlighting the importance of using appropriate statistical tests to control this source of systematic error, rather than applying arbitrary heuristic approaches^53^. Building informative matrices from gene sets on either side of this significance threshold, and again applying both CAT+GTR mixture models and ML profile mixtures, we see strong support for Cnidaria+Placozoa in the test-passing supermatrix, and conversely, strong support for Cnidaria+Bilateria in the test-failing supermatrix (Figure 4, Figure 4 – Supplemental Figure 1, Figure 4 – Supplemental Figure 2). Interestingly, in trees built from the test-failing supermatrix (Figure 4 A,C; Figure 4 – Supplemental Figure 1), in both amino-acid and Dayhoff-6 alphabets, we also observe full support for Porifera as sister to all other animals. Indeed, Dayhoff-6 recoding appears significant only for the test-passing supermatrix (Figure 4 B,D), where it obviates support for Ctenophora-sister (Figure 4B, Figure 4 – Supplemental Figure 2) in favour of (albeit, with marginal support) Porifera-sister (Figure 4D), and also diminishes support for Placozoa+Cnidaria, perhaps reflecting the inherent information loss of using a reduced amino-acid alphabet for this relatively shorter matrix. In light of these observations, we question the assertion that the support for Porifera-sister seen in some matrices and recoding strategies can be *per se* attributed to a lessened influence of compositional heterogeneity^31^.

**Figure 4.**
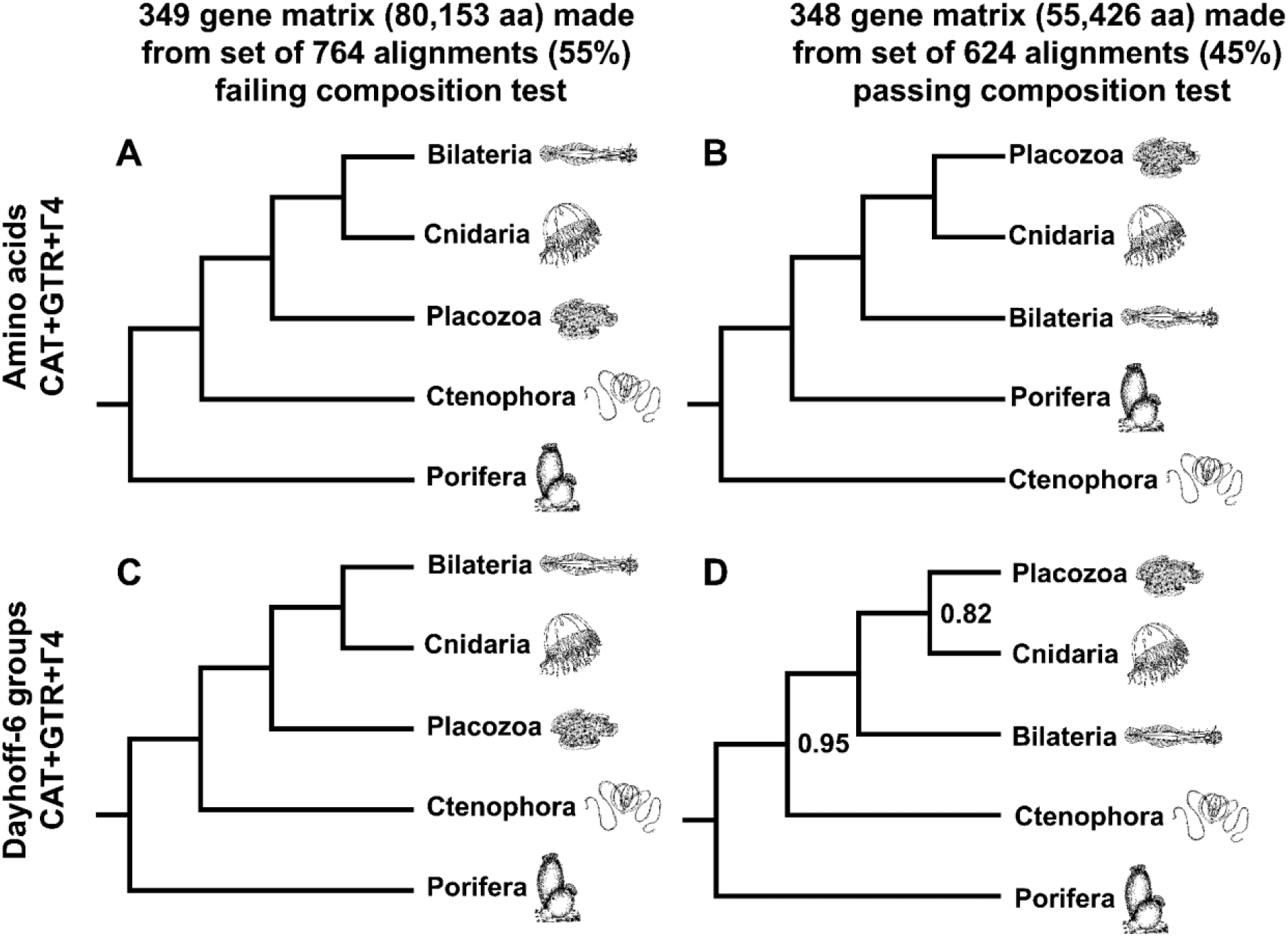
– Schematic depiction of deep metazoan interrelationships in posterior consensus trees from CAT+GTR+Г4 mixture model analyses of matrices made from subsets of genes passing or failing a sensitive null-simulation test of compositional heterogeneity. Panels correspond to A.) the amino acid matrix made within the failing set; B.) the amino acid matrix derived from the passing set; C.) the Dayhoff-6 recoded matrix from the failing set; D.) the Dayhoff-6 recoded matrix from the passing set. Only nodes with posterior probability less than 1.00 are annotated numerically.

The previously cryptic phylogenetic link between cnidarians and placozoans seen in gene sets less influenced by compositional bias will continue to raise questions on the homology of certain traits across non-bilaterians. Many workers, citing the incompletely known development^10, 12^ and relatively bilaterian-like gene content of placozoans^9, 51^, presume that these organisms must have a still-unobserved, more typical development and life cycle, or else are merely oddities that have experienced wholesale secondary simplification, having scant significance to any evolutionary path outside their own. Indeed, it is tempting to interpret this new phylogenetic position as further bolstering such hypotheses, as much work on cnidarian models in the evo-devo paradigm is predicated on the notion that cnidarians and bilaterians share, more or less, many homologous morphological features, viz. bilaterality^54^, nervous systems^36, 37, 55–57^, basement-membrane lined epithelia^58, 59^, musculature^39^, embryonic germ-layer organisation^60^, and internal digestion^38, 61–63^. While we do not argue, as some have done^64, 65^, that placozoans resemble hypothetical metazoan ancestors, we hesitate to dismiss them *a priori* as irrelevant to understanding early bilaterian evolution in particular: although apparently simpler and less diverse, placozoans nonetheless have equal status to cnidarians as an immediate extant outgroup. Rather, we see value in testing assumed hypotheses of homology, character by character, by extending pairwise comparisons between bilaterians and cnidarians to include placozoans, an agenda which demands reducing the large disparity in embryological, physiological, and molecular genetic knowledge between these taxa. Conversely, we emphasize another implication of this phylogeny: characters that can be validated as homologous at any level between Bilateria and Cnidaria must have originated earlier in animal evolution than previously appreciated, and should either cryptically occur in modern placozoans or else have been lost at some point in their ancestry. In this light, paleobiological scenarios of early animal evolution founded on inherently phylogenetically-informed interpretations of Ediacaran fossil forms^66–70^ and molecular clock estimates^71–73^ may require re-examination.

## Materials and Methods

### Sampling, sequencing, and assembling reference genomes from previously unsampled placozoans

Haplotype H4 and H6 placozoans were collected from water tables at the Kewalo Marine Laboratory, University of Hawaii-Manoa, Honolulu, Hawaii in October 2016. Haplotype H11 placozoans were collected from the Mediterranean ‘*Anthias’* show tank in the Palma de Mallorca Aquarium, Mallorca, Spain in June 2016. All placozoans were sampled by placing glass slides suspended freely or mounted in cut-open plastic slide holders into the tanks for 10 days^12^. Placozoans were identified under a dissection microscope and single individuals were transferred to 500 µl of RNA*later*, stored as per manufacturer’s recommendations.

DNA was extracted from 3 individuals of haplotype H11 and 5 individuals of haplotype H6 using the DNeasy Blood & Tissue Kit (Qiagen, Hilden, Germany). DNA and RNA from three haplotype H4 individuals were extracted using the AllPrep DNA/RNA Micro Kit (Qiagen), with both kits used according to manufacturer’s protocols.

Illumina library preparation and sequencing was performed by the Max Planck Genome Centre, Cologne, Germany. In brief, DNA/RNA quality was assessed with the Agilent 2100 Bioanalyzer (Agilent, Santa Clara, USA) and the genomic DNA was fragmented to an average fragment size of 500 bp. For the DNA samples, the concentration was increased (MinElute PCR purification kit; Qiagen, Hilden, Germany) and an Illumina-compatible library was prepared using the Ovation® Ultralow Library Systems kit (NuGEN, Leek, The Netherlands) according the manufacturer’s protocol. For the haplotype H4 RNA samples, the Ovation RNA-seq System V2 (NuGen, 376 San Carlos, CA, USA) was used to synthesize cDNA and sequencing libraries were then generated with the DNA library prep kit for Illumina (BioLABS, Frankfurt am Main, Germany). All libraries were size selected by agarose gel electrophoresis, and the recovered fragments quality assessed and quantified by fluorometry. For each DNA library 14-75 million 100 bp or 150 bp paired-end reads were sequenced on Illumina HiSeq 2500 or 4000 machines (Illumina, San Diego, U.S.A); for the haplotype H4 RNA libraries 32-37 million single 150 bp reads were obtained.

For assembly, adapters and low-quality reads were removed with bbduk (https://sourceforge.net/projects/bbmap/) with a minimum quality value of two and a minimum length of 36 and single reads were excluded from the analysis. Each library was error corrected using BayesHammer^74^. A combined assembly of all libraries for each haplotype was performed using SPAdes 3.62^75^. Haplotype 4 and H11 data were assembled from the full read set with standard parameters and kmers 21, 33, 55, 77, 99. The Haplotype H6 data was preprocessed to remove all reads with an average kmer coverage <5 using bbnorm and then assembled with kmers 21, 33, 55 and 77.

Reads from each library were mapped back to the assembled scaffolds using bbmap (https://sourceforge.net/projects/bbmap/) with the option fast=t. Scaffolds were binned based on the mapped read data using MetaBAT^76^ with default settings and the ensemble binning option activated (switch –B 20). The *Trichoplax* host bins were evaluated using metawatt^77^ based on coding density and sequence similarity to the *Trichoplax* H1 reference assembly (<u>NZ_ABGP00000000.1</u>). The bin quality metrics were computed with BUSCO2^52^ and QUAST^78^.

### Predicting proteomes from transcriptome and genome assemblies

Predicted proteomes from species with published draft genome assemblies were downloaded from the NCBI Genome portal or Ensembl Metazoa in June 2017. For Clade A placozoans, host metagenomic bins were used directly for gene annotation. For the H6 and H11 representatives, annotation was entirely *ab initio*, performed with GeneMark-ES^79^; for the H4 representative, total RNA-seq libraries obtained from three separate isolates (SRA accessions XXXX, XXXX, and XXXX) were mapped to genomic contigs with STAR v2.5.3a^80^ under default settings; merged bam files were then used to annotate genomic contigs and derive predicted peptides with BRAKER v1.9^81^ under default settings. Choanoflagellate proteome predictions^30^ were provided as unpublished data from Dan Richter. Peptides from a *Calvadosia* (previously *Leucosolenia*) *complicata* transcriptome assembly were downloaded from <u>compagen.org</u>. Peptide predictions from *Nemertoderma westbladi* and *Xenoturbella bocki* as used in Cannon et al 2016^82^ were provided directly by the authors. The transcriptome assembly (raw reads unpublished) from *Euplectella aspergillum* was provided by the Satoh group, downloaded from (http://marinegenomics.oist.jp/kairou/viewer/info?project_id=62). Predicted peptides were derived from Trinity RNA-seq assemblies (multiple versions released 2012-2016) as described by Laumer et al.^83^ for the following sources/SRA accessions:: Porifera: *Petrosia ficiformis*: SRR504688, *Cliona varians*: SRR1391011, *Crella elegans*: SRR648558, *Corticium candelabrum*: SRR504694-SRR499820-SRR499817, *Spongilla lacustris*: SRR1168575, *Clathrina coriacea*: SRR3417192, *Sycon coactum*: SRR504689-SRR504690, *Sycon ciliatum*: ERR466762, *Ircinia fasciculata*, *Chondrilla caribensis* (originally misidentified as *Chondrilla nucula*) and *Pseudospongosorites suberitoides* from (https://dataverse.harvard.edu/dataverse/spotranscriptomes); Cnidaria: *Abylopsis tetragona*: SRR871525, *Stomolophus meleagris*: SRR1168418, *Craspedacusta sowerbyi*: SRR923472, *Gorgonia ventalina*: SRR935083; Ctenophora: *Vallicula multiformis*: SRR786489, *Pleurobrachia bachei*: SRR777663, *Beroe abyssicola*: SRR777787; Bilateria: *Limnognathia maerski*: SRR2131287. All other peptide predictions were derived through transcriptome assembly as paired-end, unstranded libraries with Trinity v2.4.0^84^, running with the –trimmomatic flag enabled (and all other parameters as default), with peptide extraction from assembled transcripts using TransDecoder v4.0.1 with default settings. For these species, no ad hoc isoform selection was performed: any redundant isoforms were removed during tree pruning in the orthologue determination pipeline (see below).

### Orthologue identification and alignment

Predicted proteomes were grouped into top-level orthogroups with OrthoFinder v1.0.6^85^, run as a 200-threaded job, directed to stop after orthogroup assignment, and print grouped, unaligned sequences as FASTA files with the ‘-os’ flag. A custom python script (‘renamer.py’) was used to rename all headers in each orthogroup FASTA file in the convention [taxon abbreviation] + ‘@’ + [sequence number as assigned by OrthoFinder SequenceIDs.txt file], and to select only those orthogroups with membership comprising at least one of all five major metazoan clades plus outgroups, of which exactly 4,300 of an initial 46,895 were retained. Scripts in the Phylogenomic Dataset Construction pipeline^86^ were used for successive data grooming stages as follows: Gene trees for top-level orthogroups were derived by calling the fasta_to_tree.py script as a job array, without bootstrap replicates; six very large orthogroups did not finish this process. In the same directory, the trim_tips.py, mask_tips_by_taxonID_transcripts.py, and cut_long_internal_branches.py scripts were called in succession, with ‘./ .tre 10 10’, ‘./ ./ y’, and ‘./ .mm 1 20 ./’ passed as arguments, respectively. The 4,267 subtrees generated through this process were concatenated into a single Newick file and 1,419 orthologues were extracted with UPhO^87^.

Orthologue alignment was performed using the MAFFT v7.271 ‘E-INS-i’ algorithm, and probabilistic masking scores were assigned with ZORRO^88^, removing all sites in each alignment with scores below 5 as described previously^83^. 31 orthologues with retained lengths less than 50 amino acids were discarded, leaving 1,388 well-aligned orthologues.

### Matrix assembly

A full concatenation of all retained 1,388 orthogroups was performed with the ‘geneStitcher.py’ script distributed with UPhO available at https://github.com/ballesterus/PhyloUtensils. However, such a matrix would be too large for tractably inferring a phylogeny under well-fitting mixture models such as CAT+GTR; therefore we used MARE v0.1.2^40^ to extract an informative subset of genes using tree-likeness scores, running with ‘-t 100’ to retain all taxa and using ‘-d 1’ as a tuning parameter on alignment length. This yielded our 430-orthologue, 73,547 site matrix.

As a check on the above procedure, which is agnostic to the identity of the genes assigned into orthologue groups, we also sought to construct a matrix using complete, single-copy sequences identified by the BUSCO v3.0.1 algorithm^52^, using the 303-gene eukaryote_odb9 orthologue set. BUSCO was run independently on each peptide FASTA file used as input to OrthoFinder, and a custom python script (‘extract.py’) was used to parse the full output table from each species, selecting only those entries identified as complete-length, single-copy representatives of each BUSCO orthologue, and grouping these into unix directories, facilitating downstream alignment, probabilistic masking, and concatenation, as described for the OrthoFinder matrix. This 303-gene BUSCO matrix had a total length of 94,444 amino acids, with 39.6% of sites representing gaps or missing data.

Within the gene bins nominated by the test of compositional heterogeneity (see below), matrices were constructed again by concatenating and reducing matrices with MARE, using ‘-t 100’ to retain all taxa and setting ‘-d 0.5’ to yield a matrix of an optimal size for inferring a phylogeny under the CAT+GTR model. This procedure gave a 349-gene matrix of 80,153 amino acids within the test-failing gene set, and a 348-gene matrix of 55,426 amino acids within the test-passing set (Figure 4).

### Phylogenetic Inference

Individual ML gene trees were constructed on all 1,388 orthologues in IQ-tree v1.6beta, with ‘-m MFP -b 100’ passed as parameters to perform automatic model selection and 100 standard nonparametric bootstraps on each gene tree.

For inference on the initial 430-gene matrix, we proceeded as follows: ML inference on the concatenated matrix (Figure 1 – Supplemental Figure 1) was performed with IQ-tree v1.6beta, passing ‘-m C60+LG+FO+R4 -bb 1000’ as parameters to specify a profile mixture model and retain 1000 trees for ultrafast bootstrapping; the ‘-bnni’ flag was used to incorporate NNI correction during UF bootstrapping, an approach shown to control misleading inflated support arising from model misspecification^89^. ML inference using only the H1 haplotype as a representative of Placozoa (Figure 1 – Supplemental Figure 2) was undertaken similarly, albeit using a marginally less complex profile mixture model (C20+LG+FO+R4). Bayesian inference under the CAT+GTR+Г4 model was performed in PhyloBayes MPI v1.6j ^47^ with 20 cores each dedicated to 4 separate chains, run for 2885-3222 generations with the ‘-dc’ flag applied to remove constant sites from the analysis, and using a starting tree derived from the FastTree2 program^90^. The two chains used to generate the posterior consensus tree summarized in Figure 1 converged on exactly the same tree in all MCMC samples after removing the first 2000 generations as burn-in. Analysis of Dayhoff-6-state recoded matrices in CAT+GTR+Г4 was performed with the serial PhyloBayes program v4.1c, with ‘-dc -recode dayhoff6’ passed as flags. Six chains on the 430-gene matrix were run from 1441-1995 generations; two chains showed a maximum bipartition discrepancy (maxdiff) of 0.042 after removing the first 1000 generations as burn-in (Figure 2). QuartetScores^42^ was used to measure internode certainty metrics including the reported EQP-IC, using the 430 gene trees from those orthologues used to derive the matrix as evaluation trees, and using the amino-acid CAT+GTR+Г4 tree as the reference to be annotated (Figure 1).

For inference on the BUSCO 303 gene set, we ran 4 chains of the CAT+GTR+Г4 mixture model with PhyloBayes MPI v1.7a, applying the -dc flag again to remove constant sites, but here not specifying a starting tree; chains were run from 1873 to 2361 generations. Unfortunately, no pair of chains reached strict convergence on the amino-acid version of this matrix (with all pairs showing a maxdiff = 1 at every burn-in proportion examined), perhaps indicating problems mixing among the four chains we ran. However, all chains showed full posterior support for identical relationships among the 5 major animal groups, with differences among chains assignable to minor differences in the internal relationships within Choanoflagellata and Bilateria. Accordingly, the posterior consensus tree in Figure 3A is summarized from all four chains, with a burn-in of 1000 generations, sampling every 10 generations. For the Dayhoff-recoded version of this matrix, we ran 6 separate chains again with CAT+GTR+Г4 with the -dc flag, for 5433-6010 generations; two chains were judged to have converged, giving a maxdiff of 0.141157 during posterior consensus summary with a burn-in of 2500, sampling every 10 generations (Figure 3B).

For inference on the 348 and 349 gene matrices produced within gene bins defined by the null-simulation test of compositional bias (see below), we ran 6 chains each for the amino acid and recoded versions of each matrix, under CAT+GTR+Г4 with constant sites removed. In the amino-acid matrix, chains ran from 2709-3457 and 1423-1475 generations for the test-failing and test-passing matrices, respectively. In the recoded matrix, chains ran from 3893-4480 and 4350-4812 generations for the test-failing and test-passing matrices, respectively. In selecting chains to input for posterior consensus summary tree presentation (Figures 4A-D), we chose pairs of chains and burn-ins that yielded the lowest possible maxdiff values (all <0.1 with the first 500 generations discarded as burn-in, except for the amino-acid coded test-failing matrix, whose most similar pair of chains gave a maxdiff of 0.202 with 1000 generations discarded as burn-in). We emphasize that the topologies and supports displayed in Figures 4A-D are similar when all chains (and conservative burn-in values) are used to generate consensus trees. For ML trees using profile mixture models for the test-failing (Figure 4 – Supplemental Figure 1) and test-passing (Figure 4 – Supplemental Figure 2) gene matrices, we used IQ-tree 1.6rc, calling in the same manner (with C60+LG+FO+R4) as used on our 430-gene matrix (see above).

### Tests of compositional heterogeneity

For posterior predictive tests of compositional heterogeneity using MCMC samples under CAT+GTR, we used PhyloBayes MPI v1.7a to test two chains from the initial 430-gene matrix, and 3 chains from the 303-gene BUSCO matrix, removing 2000 and 1000 generations as burn-in, respectively. Results from tests on representative chains were selected for plotting in Figure 3C; however, results from all chains tested are deposited in the Data Dryad accession.

For the per-gene null simulation tests of compositional bias^45^, we used the p4 package (https://github.com/pgfoster/p4-phylogenetics), inputting the ML trees inferred by IQ-tree for each of the 1,388 alignments, and assuming an LG+Γ4 substitution model with a single empirical frequency vector for each gene; this test was implemented with a simple wrapper script (‘p4_compo_test_multiproc.py’) leveraging the python multiprocessing module. We opted not to model-test each gene individually in p4, both because the range of models implemented in p4 are more limited than those tested for in IQ-tree, and because, as a practical matter, LG (usually with variant of the FreeRates model of rate heterogeneity) was chosen as the best-fitting model in the IQ-tree model tests for a large majority of genes, suggesting that LG+Γ4 would be a reasonable approximation for the purposes of this test. We selected an α-threshold of 0.10 for dividing genes into test-passing and -failing bins as a conservative measure; however, we emphasize that even at a less conservative α=0.05, 47% of genes would still be detected as falling outside the null expectation.

## Acknowledgements

Nicole Dubilier (Max Planck Institute for Marine Microbiology) contributed resources that permitted the collection and assembly of draft *Trichoplax* genomes, which were amplified and sequenced at the Max Planck-Genome-Centre Cologne. Dan Richter (King lab) and Kanako Hisata (Satoh lab) provided access to unpublished transcriptomes and peptide predictions. The EMBL-EBI Systems Infrastructure team provided essential support on the EBI compute cluster. Allen Collins, Scott Nichols, and particularly Andreas Hejnol provided useful comments on an earlier version of this manuscript.

## Source data availability

SRA accession codes, where used, and all alternative sources for sequence data (e.g. individually hosted websites, personal communications), are listed above in the Materials and Methods section. A DataDryad accession is available at https://doi.org/10.5061/dryad.6cm1166, which makes available all helper scripts, orthogroups, multiple sequence alignments, phylogenetic program output, and raw host proteomes inputted to OrthoFinder. Metagenomic bins containing placozoan host contigs and raw RNA reads used to derive gene annotations from H4, H6 and H11 isolates are also provided in this accession. PhyloBayes .chain files, due to their large size, are separately accessioned at in Zenodo at https://doi.org/10.5281/zenodo.1197272.

## Author Contributions

CEL assembled most libraries, conducted all analyses starting from predicted peptides onwards, and wrote the initial draft. HGV collected clade A placozoan isolates from Majorca, maintained Hawaiian isolates prior to sequencing at the MPI for Marine Microbiology, submitted purified nucleic acids for amplification and sequencing, and assembled and provided binned metagenomic contigs. MGH and VBP assisted with collection of the original Hawaiian placozoan isolates, and H4 and H6 samples used for sequencing were derived from clones originally established by MGH at the Kewalo Marine Labs. AR generated new transcriptomic data for many sponge taxa. CEL, JCM and GG conceptualized and initiated this work, and supervised throughout. All authors read and contributed to the final manuscript.

---

**Figure 1 – Supplementary Figure 1.**
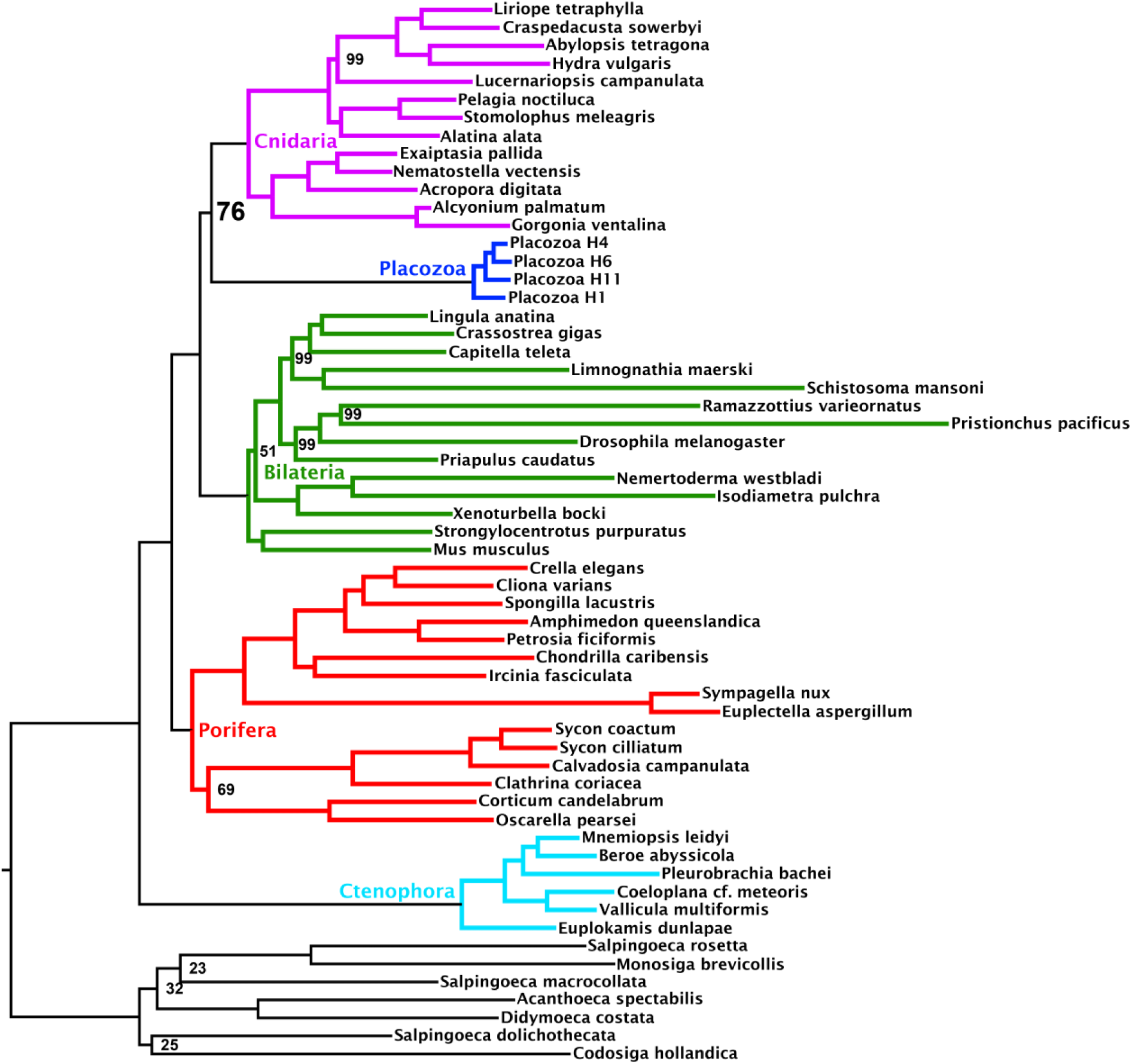
Maximum likelihood tree under the C60+LG+FO+R4 profile mixture model, inferred from the 430-orthologue matrix with full taxon sampling. Nodes annotated with ultrafast bootstrap supports with NNI correction; unannotated nodes received full support.

**Figure 1 – Supplementary Figure 2.**
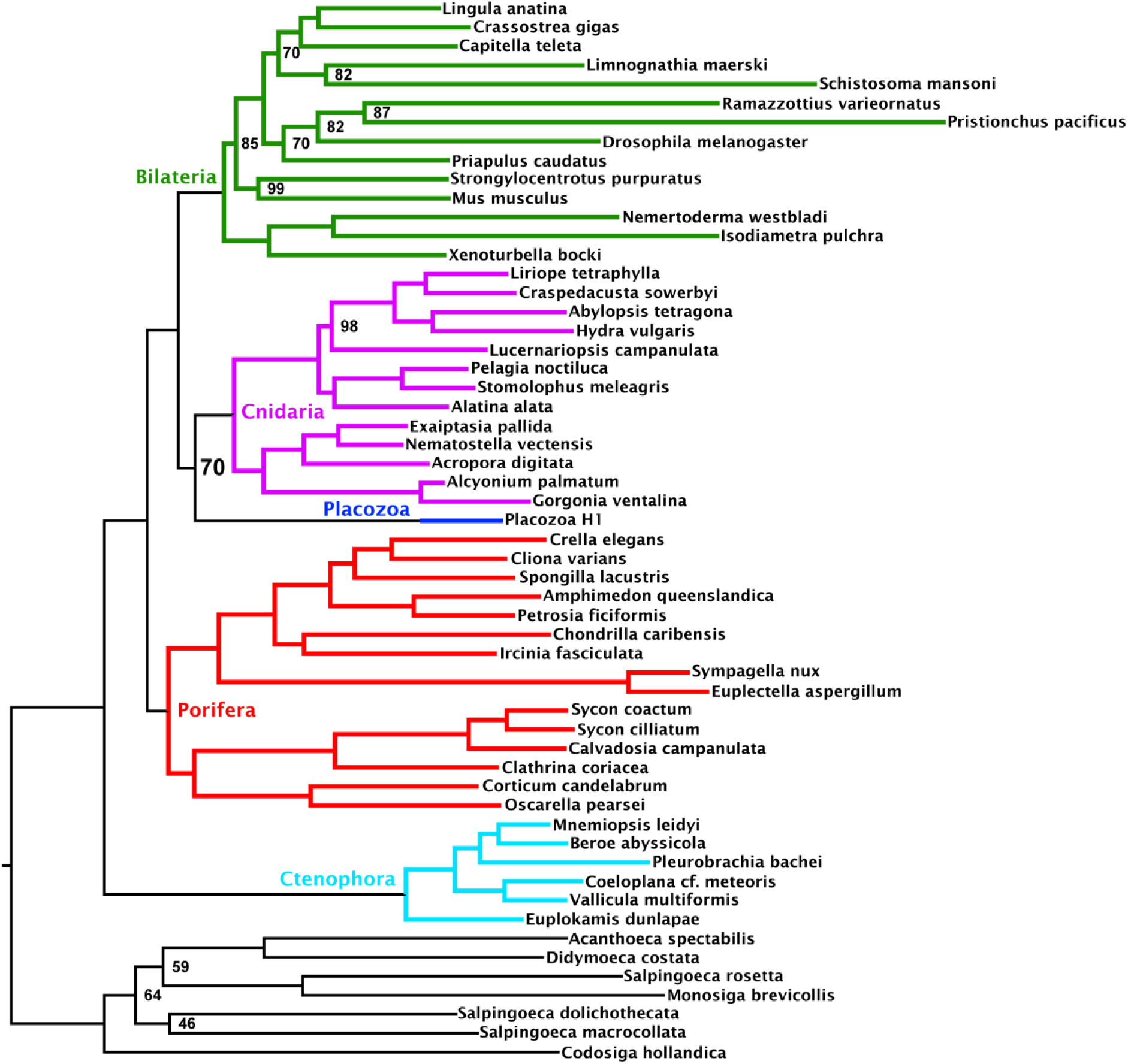
Maximum likelihood tree under a profile mixture model inferred from the 430-orthologue matrix, with only Placozoa H1 used to represent this clade. Nodes annotated with ultrafast bootstrap supports with NNI correction; unannotated nodes received full support.

**Figure 4 – Supplementary Figure 1.**
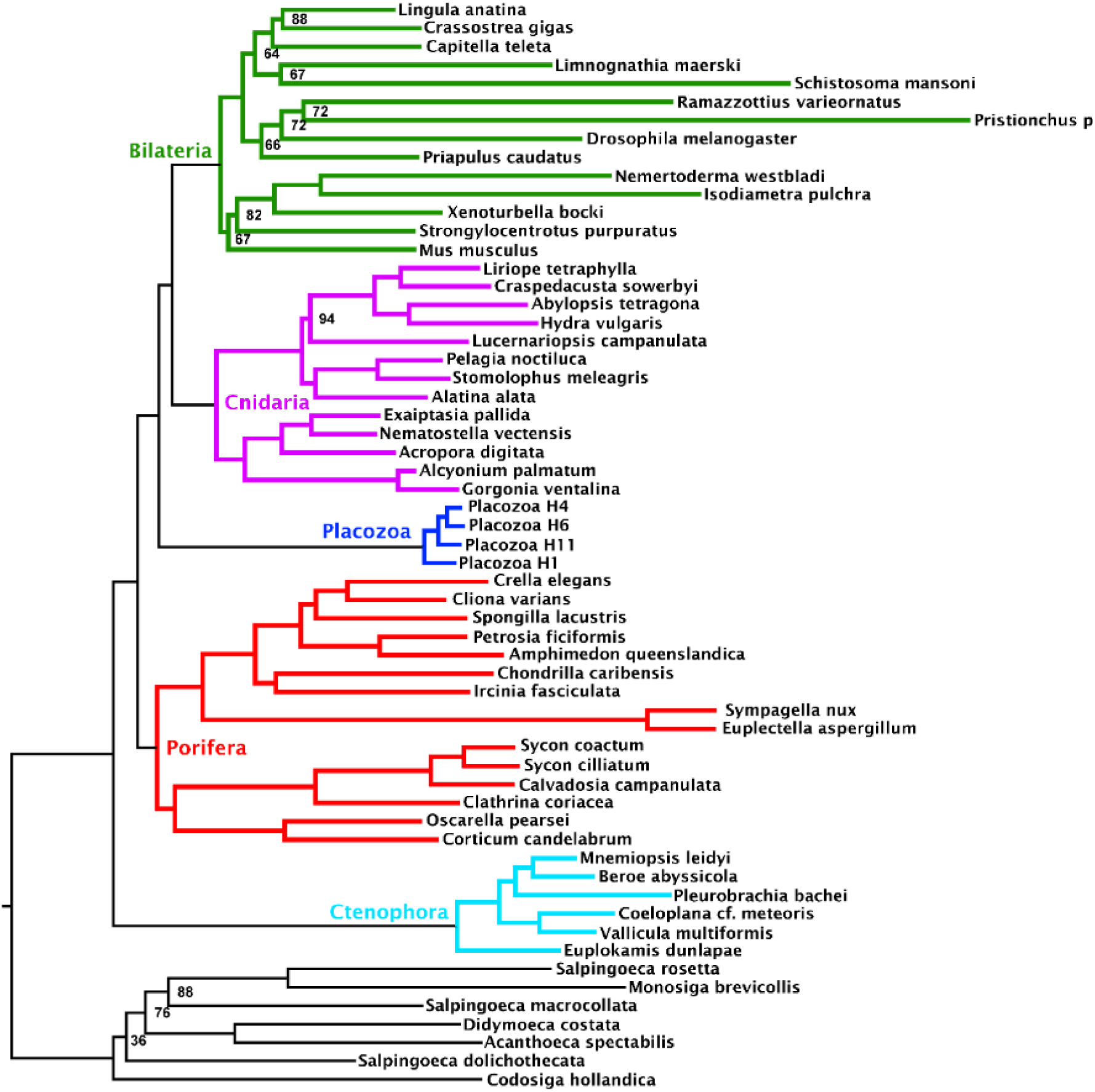
Maximum likelihood tree under a profile mixture model inferred from the 349-orthologue matrix composed from the subset of genes binned as failing the null-simulation compositional bias test. Nodes annotated with ultrafast bootstrap supports with NNI correction; unannotated nodes received full support.

**Figure 4 – Supplementary Figure 2.**
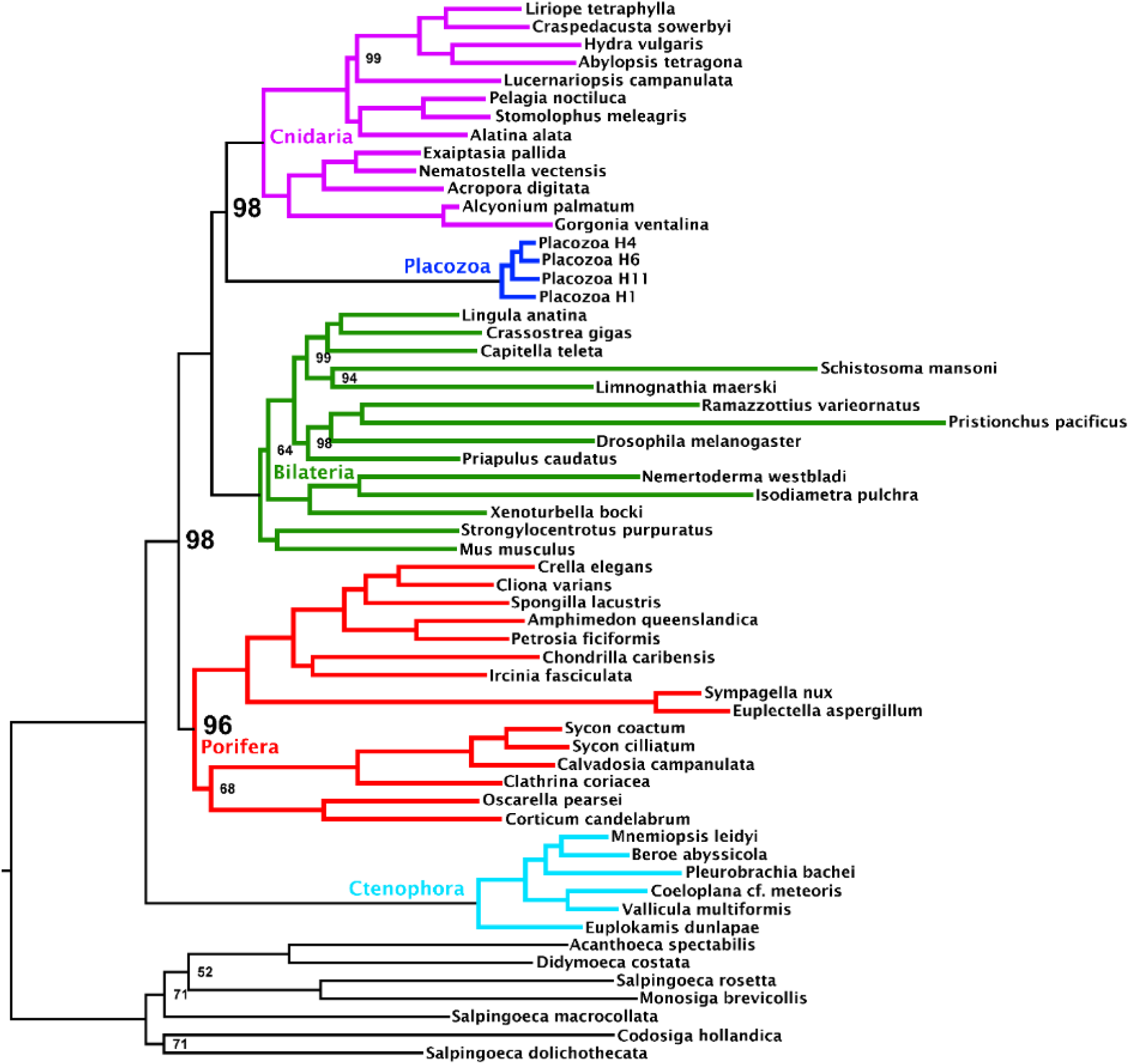
Maximum likelihood tree under a profile mixture model inferred from the 348-orthologue matrix composed from the subset of genes binned as passing the null-simulation compositional bias test. Nodes annotated with ultrafast bootstrap supports with NNI correction; unannotated nodes received full support.

